# Perception and neural representation of intermittent odor stimuli in mice

**DOI:** 10.1101/2025.02.12.637969

**Authors:** Luis Boero, Hao Wu, Joseph D. Zak, Paul Masset, Farhad Pashakhanloo, Siddharth Jayakumar, Bahareh Tolooshams, Demba Ba, Venkatesh N. Murthy

**Author notes:** Correspondence to Venkatesh N. Murthy. These authors contributed equally (Luis Boero, Hao Wu).

## Abstract

Odor cues in nature are sparse and highly fluctuating due to turbulent transport. To investigate how animals perceive these intermittent cues, we developed a behavioral task in which head-restrained mice made binary decisions based on the total number of discrete odor pulses presented stochastically over several seconds. Mice readily learned this task, and their performance was well-described by widely used decision models. Logistic regression of binary choices against the timing of odor pulses within the respiratory cycle revealed that mice placed higher perceptual weight to stimuli arriving during inhalation than exhalation, a phase dependency that strongly correlated with the magnitude of responses in olfactory sensory neurons. The population response of anterior piriform cortex (APCx) neurons to odor pulses was also modulated by respiration phase, although individual neurons displayed varying levels of phase-dependence. Single APCx neurons responded stochastically and transiently to odor pulses, leading to a representation that carries signatures of sensory evidence, but not its accumulation. Our study reveals that mice can integrate intermittent odor signals across dozens of breaths, but respiratory modulation of sensory inputs imposes limits on information acquisition that cortical circuits cannot overcome to improve behavior.

## Introduction

Many animals can navigate fluctuating odor cues to locate food or mates ^1–3^. Continuous odor gradients may be accessible for microscopic organisms, but they are rarely experienced by larger animals due to the action of turbulent fluid flow ^4,5^. Instead, animals navigating airborne odor plumes may go through a spatiotemporally discontinuous olfactory landscape, characterized by short pulses of detectable odor concentrations (‘whiffs’) interspersed with time-varying blanks ^4,6–8^. In this sparse regime, it is unclear what features of olfactory stimuli animals extract to gain information about odor sources. Physical simulations have revealed how environmental factors can affect olfactory statistics at different locations within and outside the plume. In particular, whiffs and blanks frequencies and duration, as well as odor intermittency - a related quantity measuring the probability of odor signals being above a detection threshold - were found to depend on the animal’s position within the plume ^8–10^. The spatiotemporal integration of those statistical features could allow animals to make informed decisions during odor-guided navigation, similar to the accumulation of auditory and visual information for prey localization during hunting behaviors ^11,12^.

In mice, like most other mammals, snapshots of the olfactory scene are acquired by discrete sampling events contingent on the influx of air to the nasal cavity. This is exactly what happens during normal respiration, but mice can also actively control sampling further by modulating sniffing depth and frequency ^13–15^. Once in the nasal cavity, odor molecules activate olfactory sensory neurons (OSNs) in the olfactory epithelium ^16^. Inputs from different types of OSNs converge on distinct glomeruli in the olfactory bulb (OB) ^17^, where the information is processed and transmitted to mitral and tufted cells (MTCs) that ultimately relay it to multiple areas in the olfactory cortex, hippocampus, and amygdala ^18^. The anterior piriform cortex (APCx) is one of the main targets of MTCs, plays a significant role in the representation of odor identities, and it has also extensive connections with other olfactory and non-olfactory areas ^19^.

Extensive prior work has characterized how how sniffing modulates the basal activity and the time course of neuronal responses in the olfactory pathway ^20–27^. However, those studies tended to use continuous odor stimuli extending over multiple sniffs or very brief optogenetic activations. Recent work in mice has begun to characterize the representation of fluctuating olfactory stimuli and shown that mice can discriminate variations in the frequency and phase of trains of brief, square odor pulses delivered at frequencies up to 40 Hz, probably leveraging on the high temporal fidelity of OSNs and MTCs ^28,29^. Moreover, when mice were presented with more naturalistic odor plumes, neural responses in the OB were able to follow their temporal dynamics ^30,31^, with different glomeruli tuned to different intermittency levels ^32^. Nevertheless, little is known about how fast, fluctuating stimuli such as the ones found in naturalistic odor plumes are encoded and integrated across the olfactory pathway, how odor sampling interacts with this process, and whether their representation is useful for animals to make sense of the olfactory landscape.

Here, we show that trained mice can discriminate odor stimuli with different total pulse counts, and that their decisions can be modeled well using signal detection theory. Odor pulses falling on distinct phases of breathing were perceived differently by mice, and this phase-dependence was remarkably similar to sensory responses in the OSNs. APCx neural responses to odor pulses were also modulated by breathing phase, but were stochastic and transient, which hindered accumulation of evidence. Therefore, our results provide new insights about the encoding of sub-sniff, fluctuating olfactory stimuli, while opening new questions regarding how olfactory evidence accumulation can underlie odor-guided navigation.

## Results

### Odor pulse counting task

We trained water-restricted, head-fixed mice in a two-alternative forced choice (2AFC) task in which the reward side was determined by the total number of odor pulses delivered. Animals were presented with sequences of 50 ms odor pulses of 5% ethyl valerate over a time window of 5 seconds, and the side of the reward was determined by whether the total number of odor pulses presented was above or below a fixed threshold (Figure 1A-B). Photoionization detector (PID) recordings obtained directly from the mask showed that the 50 ms voltage commands to the odor valve triggered an odor signal with ∼40 ms halfwidth (Supplementary Figure 1A), confirming that our olfactometer can produce sub-sniff olfactory stimuli. Each trial of the 2AFC task (Figure 1C) started with a sound cue marking the beginning of the sampling period in which the sequence of 50 ms odor pulses was delivered to the mask. The total number of pulses delivered in each trial was drawn with equal probability from one of two truncated Poisson distributions in which the overlap is removed, leading to trials in which the total pulse count was above or below the threshold (High and Low Trials respectively). A second sound cue denoted the end of the sampling period and the initiation of the response period in which the animal had to lick to indicate its behavioral choice. The two water ports were respectively associated with high or low number of odor pulses presented during the sampling period, and this association was counterbalanced across animals. When the mouse licked the correct water port, it was rewarded with a drop of water. When the mouse made the wrong decision, no water was given, a warning buzz sound was played, and the mouse was put under a 10-second timeout as a punishment before the next trial started.

**Figure 1.**
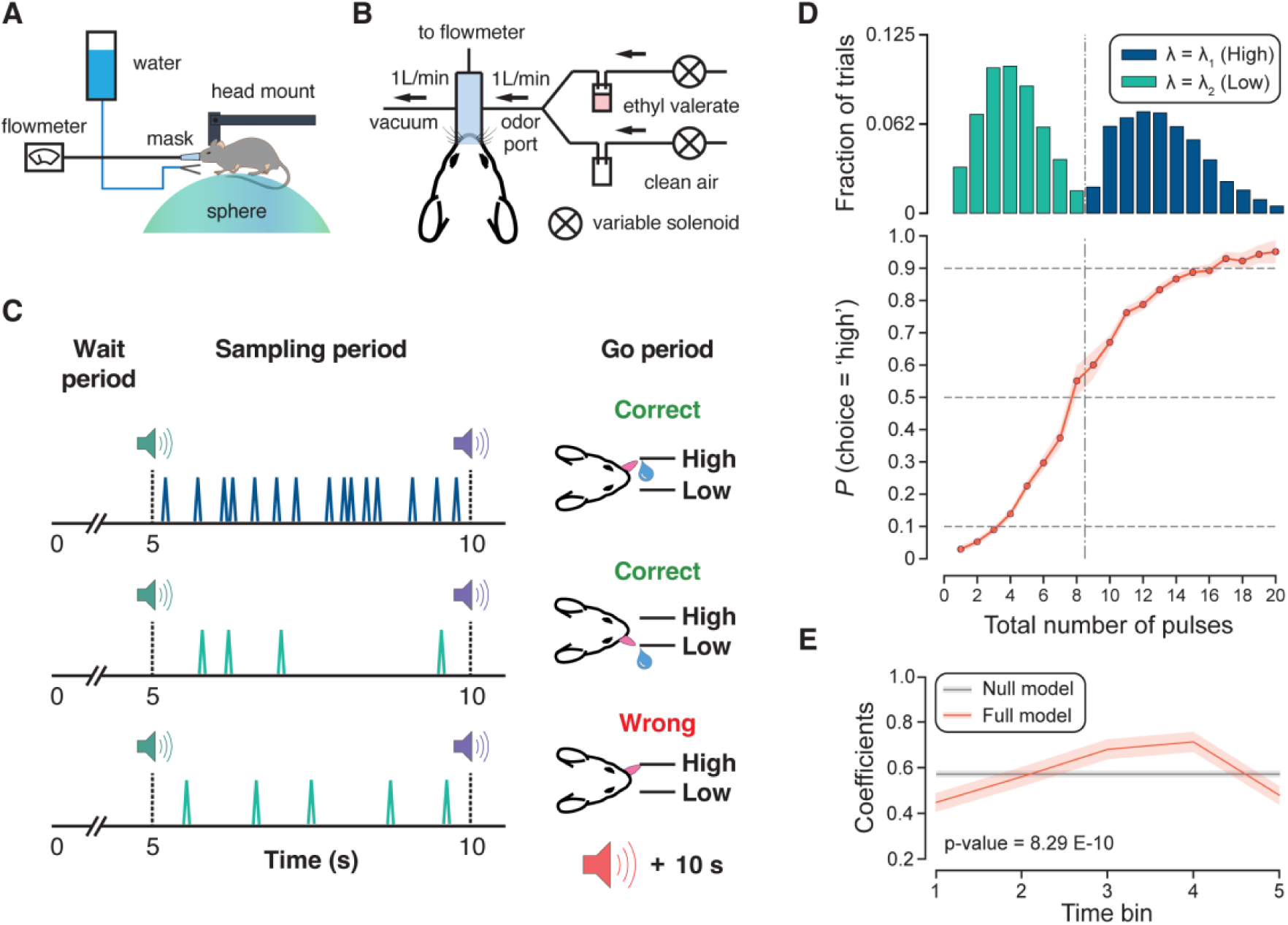
**(A)** Side view of the behavioral rig. Head-restrained mice were placed over a sphere, and a mask was fitted to their snout for odor delivery. **(B)** Schematic of the olfactometer and connections to the mask. Output of the olfactometer arrives directly to the mask, which is also connected to a vacuum line, and a flow sensor for breathing monitoring. **(C)** Schematic of the behavioral task and three example trials with their corresponding outcomes given the number of pulses delivered. **(D) Top panel**: distribution of trials collected from previously trained mice (20,042 trials across 7 mice), using a 5 s sampling window, *λ*_1_ = 12 and *λ*_2_ = 4. Gray dashed line indicates the location of the decision boundary (8.5 pulses). **Bottom panel:** psychometric curve of the pooled behavioral responses across animals under the experimental conditions described above. Orange circles indicate the mean, whereas the shade represents the 95% CI. (**E**) Mixed-effects logistic regression coefficients obtained after fitting the behavioral choices to the ‘Null’ and ‘Full’ models using odor information divided in 5 bins. Models were compared using a log-likelihood ratio test with the p-value of the comparison indicated. Data is expressed as mean ± CI 95%.

After training was completed, we analyzed the behavioral performance of mice when they were challenged with a 5 s sampling period and a ratio of 3 between the means of the two Poisson distributions used for generating the odor pulses (Figure 1D, top panel). In this context, the boundary between Low and High Trials was between 8 and 9 odor pulses. Performance approached saturation (>∼90% correct) when mice differentiated between stimuli with pulse count far from the decision boundary and degraded when pulse counts approached the boundary (Figure 1D, lower panel). Animals were also able to discriminate total pulse counts when odor pulses were delivered over shorter (1.25 s, 2.5 s) or longer (10 s) sampling windows (Supplementary Figure 1D), suggesting that mice can integrate olfactory evidence for decision-making over a wide range of intermittencies and times (Supplementary Figure 1E).

Odor pulses were presented non-uniformly and stochastically during the sampling window. Then, in principle, an accurate decision in every trial must result from an evidence accumulation process, in which the animal must integrate sensory information throughout the trial, and not just a short time window. To measure the extent to which the pulses in different parts of the trial contribute to the animal’s decision, we divided the sampling window into 5 bins (1s each) and calculated the mean odor signal for each bin in each trial. Then, binary licking decisions were fitted using two mixed effects logistic regression models: a null model in which all time bins were forced to have equal coefficients and a full model in which coefficients were allowed to differ across time bins. The full model provided a better fit to the data and was significantly different from the null model (Figure 1E, Log-likelihood ratio test, p-value: 8.29 E-10), suggesting that animals might weigh differentially sensory information coming over several seconds. Altogether, our newly developed task has allowed us to show how mice can accumulate -and weigh-odor information over several seconds for decision-making.

### Breathing influence on behavior

The sampling of olfactory inputs by mice is tied to the airflow driven by inhalation and exhalation during breathing, or its active variant, sniffing. In the context of our task, the amount of odor molecules inhaled from each pulse will depend on the arrival time of a given odor pulse relative to the breathing cycle. To test how this variability affects behavioral choice, we divided each breath into 15 bins and computed a ‘phase histogram’ representing the number of pulses that arrived at each phase bin across successive breaths on a trial (Figure 2A). When phase histograms were used as the input to train a logistic regression model to predict the licking decisions of mice in each trial, we observed that the perceptual weights (i.e., the logistic regression coefficients associated with each bin) were maximal around the peak of the inhalation and decreased progressively towards the exhalation (Figure 2B, orange trace).

**Figure 2.**
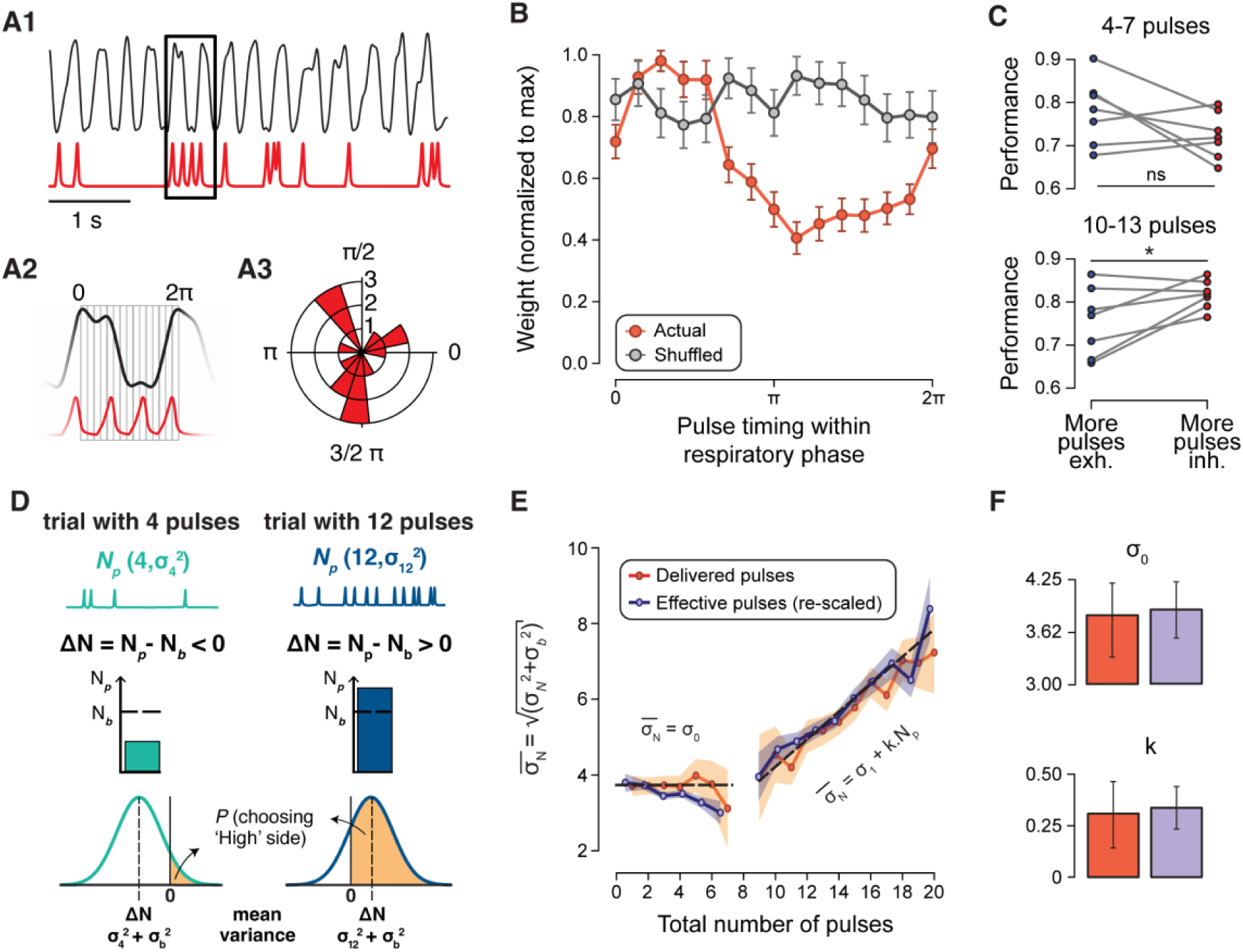
(**A1**) Breathing signal measured through the airflow sensor (upper black trace) and the odor signal delivered to the animal (lower red trace) in an example trial. (**A2**) Expansion of traces for the area delimited by the black box in A1. Each breath was divided into 15 bins from 0 to 2π to allow the sorting of odor pulses. (**A3**) Phase histogram showing the number of pulses arriving at each bin for all the breaths in the trial depicted in A1. (**B**) Normalized coefficients (weights) for each of the bins of the phase histograms used for fitting a binary logistic regression model of choices in the task. ‘Actual’ (orange trace) indicates the weights obtained based on the original phase histograms, whereas ‘Shuffled’ (gray trace) correspond to the regression weights obtained after shuffling the timing of pulses within each trial before fitting the model. Results are expressed as mean ± SD. **(C)** Comparison of within-animal performances for trials with more pulses falling during exhalation or during inhalation. Each dot corresponds to the performance of a single animal averaged across sessions for trials with 4 to 7 total pulses (left) or 10 to 13 total pulses (right). * p-value < 0.05, paired t-test). (**D**) **Top:** Schematic of two example trials with different total pulse numbers. Our hypothesis is that animals generate an estimate of the total number of pulses in each trial (*N_p_*) that follows a Gaussian distribution with a mean at the true number of pulses and a variance, σ_N_^2^, that can be different for different values of *N_p_*. **Center:** the estimate of *N_p_* will then be subtracted to the estimate of the decision boundary, *N_b_*, which we assume to also follow a Gaussian distribution with variance σ_b_^2^. **Bottom**: The difference between these two estimates (ΔN) also follows a Gaussian distribution, and then the orange shade under the positive section of the distribution represents the probability of choosing the lick port associated with the high number of pulses. **(E)** Maximum Likelihood Estimation (MLE, see Methods) of 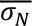 for each value of *N_p_* was computed using the delivered pulse count (orange trace) or the effective pulse count after adjusting for breathing (purple trace). 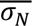 was found to be constant for low values of Np, whereas beyond the decision boundary 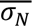 scaled with the total number of pulses presented on a trial. (**F**) σ_0_ and k values obtained from linear regression of 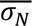 estimations based on delivered *N_p_* (orange bars) or re-scaled effective *N_p_* (purple bars). Percentile confidence intervals for σ_0 Delivered_ = (3.33, 4.21) and σ_0 Effective_ = (3.56, 4.22). Percentile confidence intervals for k_Delivered_ = (0.14, 0.46) and k_Effective_ = (0.23, 0.44).

The phase-dependence was absent when the model was trained on the same data but shuffling pulse timing during the trial (Figure 2B, gray trace). Additionally, by linearly combining the phase histogram in each trial with the mean coefficients from logistic regression, we were able to estimate the effective number of pulses experienced by mice in each trial. We found a linear relationship between the delivered pulse count and the effective pulse count, with increased variance for higher pulse counts (Supplementary Figure 2A). When effective pulse counts were binned (Supplementary Figure 2E) and used to re-compute a psychometric curve, the relationship between choice and the amount of sensory information was steeper than was observed in the original psychometric constructed using delivered pulses (Supplementary Figure 2B). These findings inspired us to classify trials based on whether more than 50% of pulses on a trial fell on the inhalation or exhalation phase (Figure 2C). We then looked at a subset of trials with total pulse counts (4-7 pulses) below the decision boundary (and found no significant differences in the performance of each mouse between the two groups of trials (paired t-test, p-value: 0.071). However, when we looked at trials with total pulse counts above the decision boundary, within-animal performance was significantly higher for trials with the majority of pulses arriving around inhalation (paired t-test, p-value: 0.021; Figure 2C). Therefore, these results indicate that the timing of fluctuating stimuli relative to the breathing cycle determines its perceptual weighing for decision-making and suggest that part of the mistakes made by animals arose from partially or totally missed pulses that arrived in proximity to exhalations.

### Models to fit behavioral data

Our experiments describe how well mice categorize odor pulses arriving in the sampling period into High or Low, and how the relative timing of a single odor pulse with respect to the animal’s respiratory cycle impacts its weighing in the context of the task. These variables not only affect how mice arrive at the estimate of the total pulse count on a trial, but also their estimation of the decision boundary used for categorizing trials as High or Low. Since both quantities are internally estimated by mice, there is uncertainty (or noise) associated with them. These uncertainties could, in principle, account for the shape of the psychometric curve, including the shallow dependence on the pulse number and the asymmetry around the decision boundary, as well as the deviation from the performance predicted by a model of a binary choices based on two different Poisson rates (Supplementary Figure 1B-C). Inspired by a previous report ^33^, we built a decision model where the perceptual estimation of the odor pulse number was Gaussian-distributed around the total number of odor pulses delivered in a trial (*N*_p_), with a variance σ^2^_N_, which could take different values for different *N*_p_ (Figure 2D). We also considered that the animal needs to generate an estimate of where the boundary between the High and Low conditions (*N_b_*) lies in the sensory space, with its corresponding error σ^2^_b_. Altogether, the ΔN value - the difference between *N*_p_ and *N*_b_ - should inform the decision of the animal: the mouse should choose the High side when ΔN > 0, and the Low side in the opposite case (Figures 2D), with an error 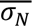 reflecting the contributions of σ^2^_N_ and σ^2^_b_. Bootstrapped estimations of 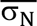 for each value of *N_p_* were generated using maximum likelihood estimation (MLE) and by setting the value of *N_b_* = 8: the pulse count closest to 0.5 in the average psychometric curve. 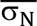 values were constant for low values of *N*_p_ but eventually started to scale with *N*_p_ (Figure 2E, orange trace).

Empirical evidence across tasks involving the simultaneous or sequential integration of quantities has supported a diverse set of models to explain the relationship between decision noise and sensory stimulus quantity ^33–35^. We found that any model assuming noise scaling with the number of pulses provided better fits to the behavioral data than a model assuming a constant noise across different numbers of pulses delivered, but the best fit to the data corresponded to a model assuming a non-zero constant noise for low pulse quantities, followed by a linear scaling of noise with higher pulse counts (Supplementary Figure 2C-D). Next, we wanted to explore whether breathing sets any constraints in the scaling of decision noise. We binned our effective pulse counts and used them to generate new estimates for 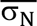 using MLE with *N_b_*= 5.01 (again, the bin center closest to 0.5 in the effective psychometric curve, Supplementary Figure 2F), and then re-scaled 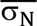 values using the regression shown in Supplementary Figure 2A. 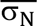 values obtained from the effective pulse counts (Figure 2E, purple trace) showed the same trend as the estimates generated with the delivered pulse count, and no significant differences were found in the values of σ_0_ and k, the parameters of the model that provided the best fit to data for the constant and linear parts of the plot, respectively (Figure 2F). This result is also supported by the fact that the psychometric curve built using effective pulse counts still shows an asymmetric dependence of choice with odor pulse counts. Taken together, our results suggest that decision noise is not constant across the whole range of total number of stimulus pulses that mice can experience in the task, and such variation is not related to sampling.

### Sensory input constrains behavior

Having discovered that the respiratory cycle governed the weighing of olfactory stimuli for the behavioral responses, we were interested in testing if that dependence was already apparent in the activity in the very first stage of odor processing, the olfactory bulb glomeruli. Therefore, we evaluated the odor pulse-triggered activity of the axonal terminals of the OSNs through wide-field calcium imaging in awake mice expressing GCaMP3 in OSNs (Figure 3A). Several glomeruli in the dorsal surface displayed detectable responses to pulses of ethyl valerate (Figure 3B), with their response timing reflecting the known kinetics of GCaMP3 ^36^, which validates the brevity of the odor pulse stimuli. The magnitude of pulse-triggered calcium signals was highly variable, even within a single glomerulus (Figure 3B; the 3^rd^, 5^th^ and 6^th^ pulses elicit negligible responses). To address if that variability in the responses was due to a breathing modulation of the sensory input, we took the same approach we used above for the behavioral data and computed the amplitude of the GCaMP3 signals as a function of pulse arrival time relative to the respiratory cycle. The average phase response curve revealed that odor pulses arriving around the peak of the inhalation phase elicited the strongest OSN calcium responses, whereas the weakest responses corresponded to pulses arriving during exhalation (Figure 3C, green trace), mirroring the dependence observed for the bins of the phase histogram for binary prediction of choices (Figure 3C, orange trace). The phase-dependence of calcium signals was observed across all the glomeruli responding to the odor pulses (Supplementary Figure 3). In fact, there was a high correlation between the logistic regression coefficients and the mean GCaMP3 pulse-triggered responses (Figure 3D, r = 0.96, p-value = 6.27E-9). Hence, the differential weighing of pulses for decision-making as a function of their arrival time in the respiratory cycle is likely constrained by the intensity of OSN responses to each pulse.

**Figure 3.**
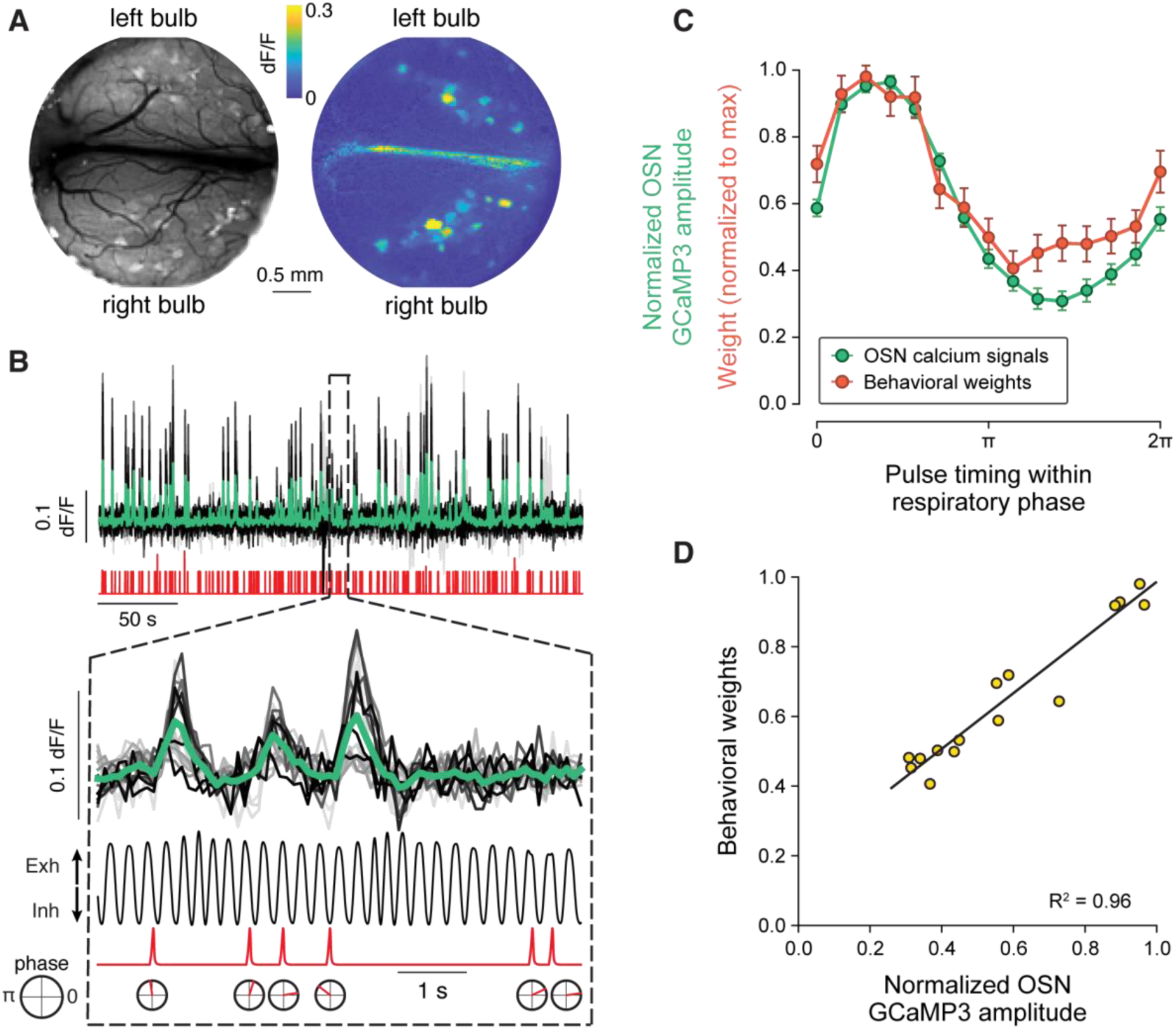
**(A)** Top view of the cranial window over the OB showing GCaMP3 signals in the OB during pulse presentation (**left**: raw image; **right**: dF/F). The ROIs with high dF/F values correspond to glomeruli responding to an odor pulse. **(B)** dF/F traces from multiple glomeruli (top trace) in response to odor pulses (bottom trace). Traces corresponding to single glomeruli are shown in grayscale, whereas the average across glomeruli is shown in green. Traces in the dashed gray box in the top panel are displayed enlarged at the bottom, with the addition of the breathing trace and a depiction of the arrival time of each of the pulses relative to the respiratory signal. **(C)** Green curve: Average pulse-triggered calcium signals (normalized to the maximum response) plotted against the phase of the breathing cycle in which the pulse arrived. Error bar corresponds to the 95% confidence interval. Amplitude of pulse-triggered calcium responses depended on the arrival time of odor pulses relative to the breathing cycle. Behavioral data from Figure 2B is reproduced here for comparison (red) and is expressed as mean ± sd. **(D)** Scatter plot of the pairs of phase bin weights for choice prediction and the normalized averaged OSN activity for each bin, and the corresponding regression line (black trace, R^2^ = 0.96, p-value = 6.51 E-9). Sample size of calcium imaging data (n = 3 animals).

### Olfactory cortical responses largely reflect sensory information

The anterior piriform cortex (APCx) is one of the main areas receiving inputs from the olfactory bulb glomeruli, and in turn it connects to the prefrontal cortex. These observations have supported the idea of APCx as a possible hub for transformation of the olfactory representations into a category-related signal interpretable for decision-making centers in the brain ^37–41^. Therefore, we were interested in assessing how the sequences of intermittent odor pulses were represented in the APCx in expert mice.

Extracellular spike recordings in APCx from trained animals during behavior revealed that the activity of many neurons was correlated with the delivery of odor pulses, exhibiting greater activity fluctuations over the 5 second period with more pulses (Figure 4A). We selected neurons whose activity was modulated by odor stimuli using a predictive criterion as described in the Methods (see *Analysis of APCx spiking data* in the Methods section). We found that many neurons exhibited graded activity (summed over 5 seconds) as pulse counts changed (Figure 4B). We first analyzed the variation in firing rates of neurons across trials with different total pulse counts (binned into 4 groups: 1-4, 5-8, 9-12, >13 pulses). We normalized the activity of a neuron in each trial to the mean activity across all trials, and classified neurons based on activity being above the mean with increasing number of pulses (57.2%), above the mean with decreasing number of pulses (39.9%), and relatively homogeneous across all pulse counts (2.9%). We then computed - for the two main groups - the average firing rates for different ranges of total pulse counts during the sampling window of the task (Figure 4C). We interpret the first group (Figure 4C, top) to have odor pulse-triggered increases in activity, and the second group (Figure 4C, bottom) to arise from odor-triggered inhibition of activity, such that these neurons spike less when there are more pulses in a trial. Importantly, we did not observe a significant cumulative increase in the firing rate over the 5 second period, as proposed for evidence accumulation areas in other studies ^42–44^.

**Figure 4.**
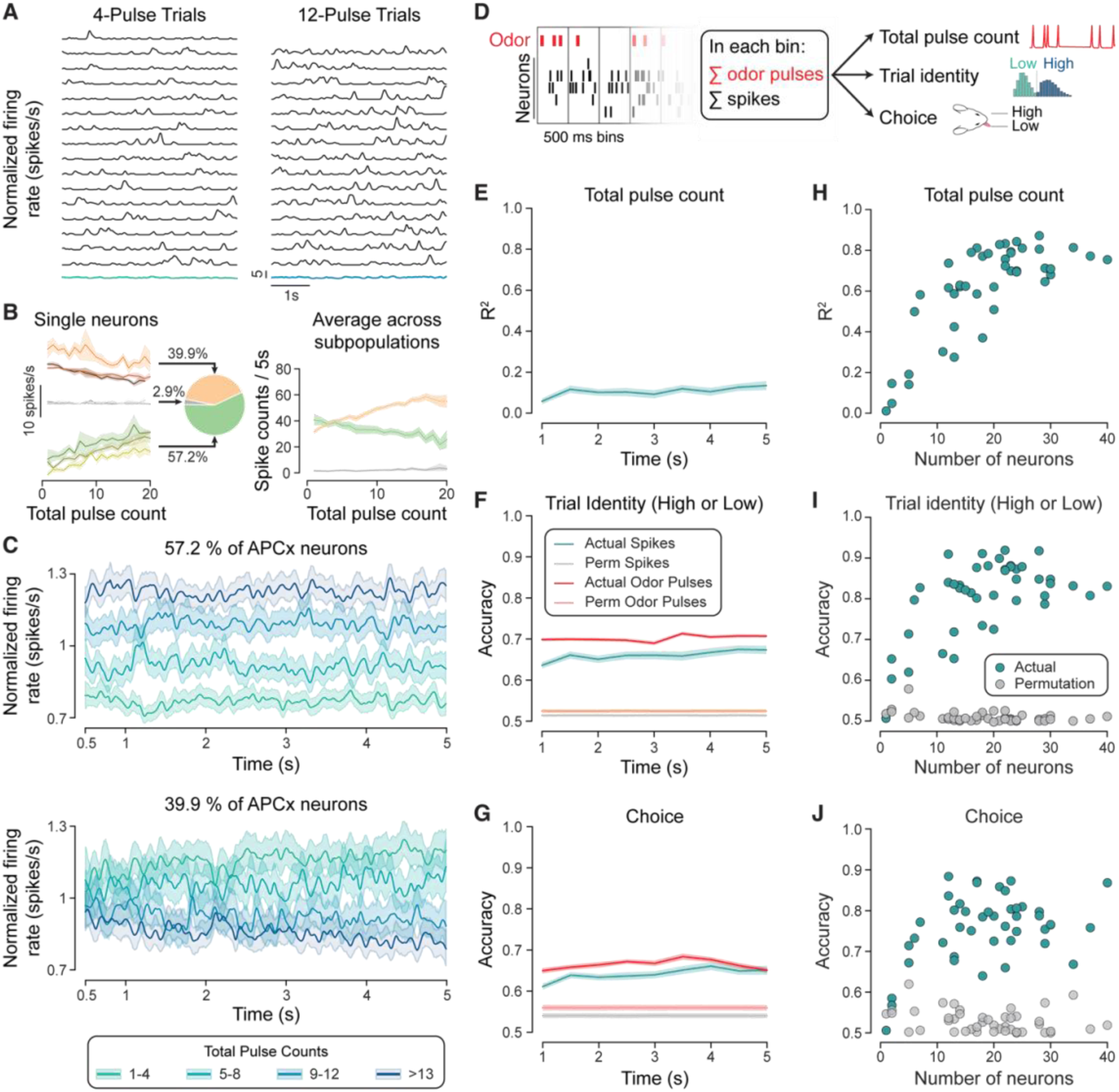
(**A**) Time course of the firing rates of a single APCx neuron across all trials in a session with 4 pulses (left) or 12 pulses (right). Colored traces in the bottom of each panel are the average response across trials. **(B) Left**: Average spike counts measured over the 5s sampling window as a function of the total number of pulses in a trial. Each coloured line represents a single neuron, and the shaded band correspond to the 95% CI. **Center**: After normalizing single neuron responses to the mean response of each neuron, APCx neurons could be classified in three main groups depending on firing rates becoming higher (green, 57.2% of APCx neurons), lower (beige, 39.9% of APCx neurons) or unaffected (grey, 2.9% of APCx neurons) with increasing number of pulses. **Right**: Average spike count for each subpopulation described before as a function of the total number of pulses in a trial. (**C**) Average firing rates of APCx neurons for ranges of total pulse counts. 57.2% of the neurons showed a higher firing rate with increasing number of pulses (**top**), whereas 39.9% of the neurons displayed firing rates above their means with decreasing number of pulses (**bottom**). **(D)** Decoding analysis rationale. Each trial was divided in 500 ms bins, and for each bin we calculated the sum of odor pulse counts as well as the summed spikes of every neuron in the recorded population. Total pulse count, trial identity (i.e. whether it was a ‘High’ or a ‘Low’ trial) and the choice of the mouse in each trial were then predicted based on the spike counts in each 500 ms bin. Prediction of trial identity and choice based on the sum of pulses in each bin was added as a measure of the maximal accuracy that can be reached based on the sensory information available at each bin. (**E**) Average multiple linear regression R^2^ using population spike counts over independent 500 ms intervals to predict total pulse counts. (**F-G**) Teal traces depict prediction accuracy of logistic regression on trial identity **(F)** and behavioral choice **(G)** using population spike counts over independent 500 ms intervals. Gray traces are the accuracies obtained when target labels were permuted. Red traces show the decoding based on the odor pulse information available, whereas pink traces correspond to the accuracies after permutation of samples. **(H)** Average R^2^ score of multiple linear regression between population spike counts and total pulse count in a trial against the number of recorded neurons in a session. (**I-J**) Same as F, but for the dependence in the accuracy of logistic regression prediction on trial identity **(J)** and animal’s decision **(J)** using population spike counts of neurons in a trial. Teal dots indicate the accuracies calculated based on the actual data, whereas gray dots are the accuracies after permutation of samples. Data is shown as mean ± CI 95%, n = 3 animals (over multiple sessions).

The overall flat structure of the average activity of neurons over the trial period of 5 seconds suggests that the information encoded by APCx neurons reflects ongoing sensory input. To test whether there was some latent accumulating information not revealed by simple trial averaging, we performed different decoding analyses to test whether the APCx activity in the late sampling period close to the time of decision making was different from earlier periods (Figure 4D). We calculated the R^2^ value of linear regression between the spike counts for each trial in successive 500 ms windows (a total 10 such bins in 5 seconds) and the total number of pulses in the trial – this regression value was low, and uniform across the entire sampling period (Figure 4E). We also used a logistic regression model to predict the trial identity (Low or High), as well as the animal’s choice using the spike counts in the successive 500ms sampling window (Figure 4F-G). Accuracy of predictions on actual data (‘Actual spikes’) was higher than for data in which labels were shuffled (‘Perm spikes’), but still below the maximal accuracy given by the actual stimulus pulse (‘Actual Odor Pulses’). Remarkably, accuracy levels of predictions made from spike counts were uniform, with no evidence that there was higher decodability closer to the decision time.

Neural activity in short time windows in APCx was not sufficient to predict task variables. We next investigated whether any downstream areas integrating information from APCx neurons over the entire 5 second sampling period will be able to recover task information. First, for each recording session, we calculated the R^2^ value of the linear regression between the total spike counts of all neurons and the total pulse count over the 5 second period in a trial. Across different sessions, this value increased with the number of recorded neurons available until a plateau was reached for population sizes higher than 20 neurons (Figure 4H). The ceiling of 20 neurons was also observed when total spike counts were fed into a logistic regression model to predict trial identity (Figure 4I) or the behavioral choice (Figure 4J). Cumulative population spike counts predicted trial identity better than choice (comparing Figure 4I and J). These data indicate that collective neural activity in a small population of APCx neurons over the entire 5 second period can be integrated downstream to make accurate binary decisions, but the activity in APCx at the end of the sampling period does not reflect decision variables.

### Detailed structure of cortical responses to fluctuating stimuli

The stochastic nature of stimulus presentation and the non-repeating structure across trials in our task precluded simple trial-averaging to reveal neuronal responses to individual odor pulses (which will occur at distinct times across trials) (Figure 5A). To overcome this limitation, we implemented a deconvolutional unrolled neural learning (DUNL) method ^45^, which uses a deep learning framework to learn locally low-rank structures from neural data. In particular, it is useful in characterizing neural responses to sparse discrete events in experiments with non-repeating trial structure or with no structure. We used DUNL to independently learn, for each neuron, its response kernel to odor pulses as well as the response intensity to each specific pulse (Figure 5B-C). Different neurons had distinct response kernels (which are essentially the impulse responses of each neuron to the odor pulse), but K-means clustering revealed the existence of 4 different groups, suggesting that the population of APCx neurons was heterogeneous in terms of how every neuron transformed the sensory input into firing (Supplementary Figure 4A). Using this method, we were also able to estimate that each odor pulse activated a subpopulation of recorded neurons, but the size and neuronal identities of those subpopulations varied from pulse to pulse. Overall, a neuron detected ∼50% of odor pulses (Figure 5D), and an odor pulse triggered the activation of ∼50% of the recorded neurons (Figure 5E). These findings clearly indicate that reliable encoding of each odor pulse must occur at the population level, rather than at the level of single neurons, in line with our analysis in Figure 4.

**Figure 5.**
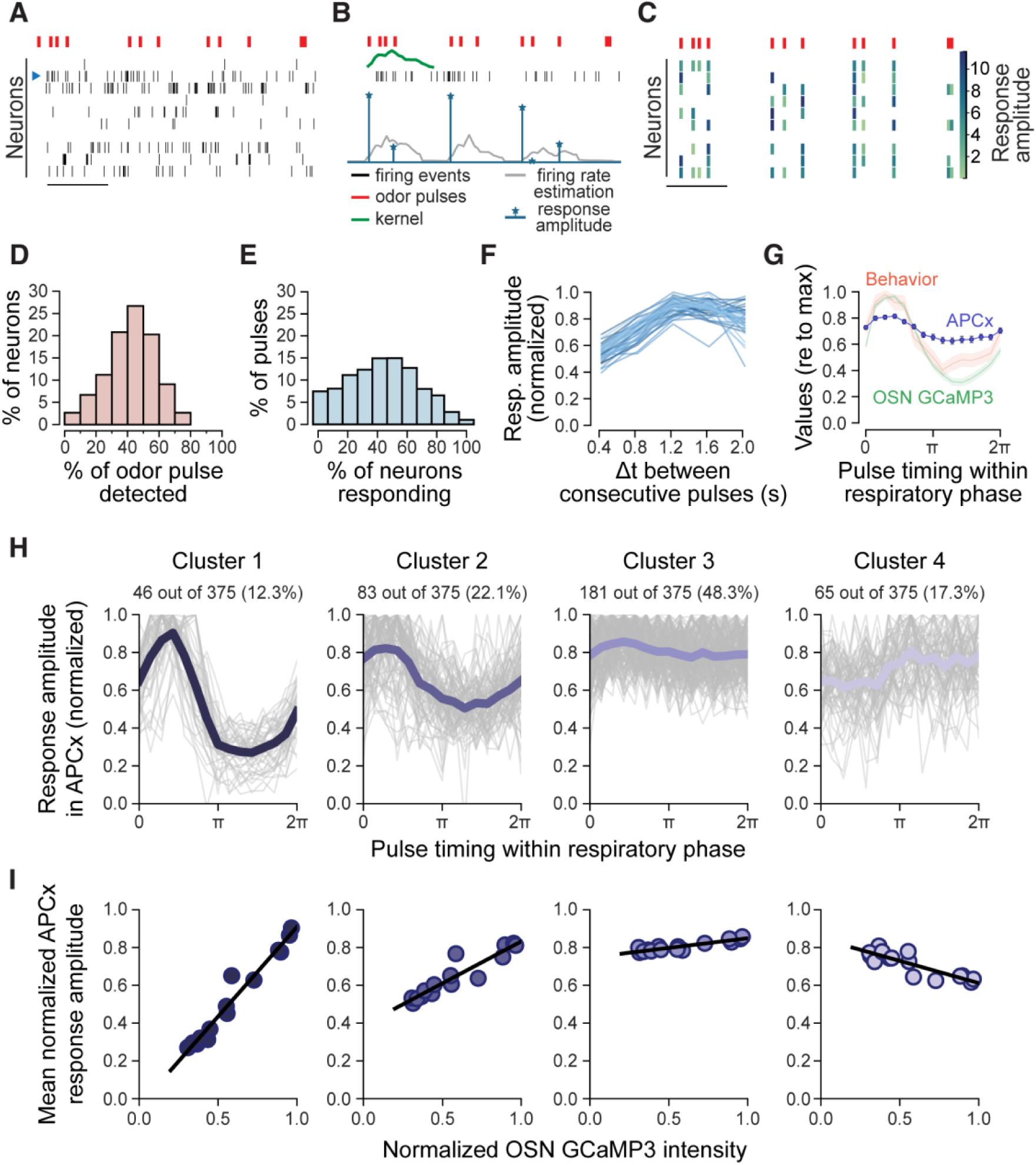
Example of the implementation of the DUNL method for the neuron marked with the blue arrowhead in the raster plot shown in (**A**) for a specific trial. (**B**) The timing of spikes and odor pulses are used to train a model that learns a specific event-based kernel for each neuron (‘kernel’, blue trace), as well as which sensory stimuli were detected and the intensity of the neuronal response (green trace, also referred as ‘response amplitude’). The convolution of response amplitudes with the kernel provides an estimation of the firing rate of the neuron (‘firing rate estimation’, orange trace). (**C**) Raster plot of the same trial of A, but with every line now showing the response amplitude in a color-coded scale. (**D**) Distribution of the percentage of neurons detecting different proportions of odor pulses presented. (**E**) Distribution of the percentage of odor pulses that elicited spiking in different proportions of the neural population. (**F**) Variation in response amplitudes as a function of the time between consecutive pulses. Each curve is the average for every neuron. **(G)** Average pooled APCx response amplitudes (mean ± CI 95%, normalized to the maximum of each neuron) against pulse timing within the respiratory cycle. Behavioral weights (mean ± sd) and calcium signals (mean ± CI 95%) in OSNs from Figure 3B (orange and green traces, respectively) are also displayed for comparison. (**H**) K-means clustering of normalized APCx neuronal responses as a function of the arrival time of odor pulses during the respiratory phase led to 4 different clusters. Gray traces indicate single-neuron responses, the blue trace shows the average of all neurons in the cluster. (**I**) Mean APCx phase responses in each cluster plotted against mean OSN responses (from Figure 3C) for each of the 15 bins of the respiratory phase. n = 3 animals over multiple sessions.

In addition to revealing which specific event any given neuron responded to, the DUNL method allowed us to infer a representation, estimating the intensity of odor response (“response amplitude” in Figure 5C), from the spiking frequency of APCx neurons. We then asked whether the intensity of the response to a given odor pulse in a single neuron was affected by the timing of the preceding odor pulse. We found that response amplitudes were reduced for the second pulse when the time between pulses was less than one second, presenting a signature of sensory adaptation in APCx neuronal firing (Figure 5F). This analysis of pairwise interactions did not consider the timing of the pulse arrival with respect to the breathing phase but averaged it over all the different arrival times.

Since olfactory sensory responses were modulated by the respiratory phase (Figure 3C), we next analyzed the relationship between cortical neuron responses and the timing of the pulses within the respiratory phase. We calculated the average response amplitude for all neurons after binning the stimuli in the same 15 bins as for behavioral and OSN calcium data. Compared to OSNs responses or the behavioral weights, the dependence of response intensity on the phase of pulse arrival was shallower, but pulses arriving around the peak of inhalation still triggered the largest responses (Figure 5G). K-means clustering of the normalized response amplitudes of individual APCx neurons over the entire respiratory phase suggested the existence of four different functional groups (Figure 5H). For comparison, we plotted the mean normalized response amplitudes in each cluster and the normalized average OSN responses from the bulbar imaging, with their corresponding linear regression (Figure 5I). One cluster -comprising ∼50% of the APCx neurons-showed no respiratory phase tuning (Figure 5H, Cluster 3), whereas the other three clusters displayed different levels of modulation of the neuronal responses by pulse timing during a breath (Figure 5H, Clusters 1,2,4). The distributions of response intensities were not significantly different across phase clusters (Supplementary Figure 4B-C), suggesting that the observed phase modulations are not due to difference in the amplitude of neuronal responses. The clustering of modulatory shapes was supported by further analysis based on individual neurons and kernel density estimation of the regression slopes (Supplementary Figure 4D-F). No correspondence was found between the clusters of neurons arising from the respiratory phase modulated responses and clusters based on the kernel shape (Supplementary Figure 4G). Therefore, fast, intermittent olfactory stimuli are represented in the APCx by neuronal ensembles that are heterogeneous in terms of their respiratory phase tuning and firing properties, features that may underlie the stochastic recruitment of neurons upon odor-pulse presentation.

Our behavioral results indicate that odor pulses arriving at different phases of respiration have different perceptual weights (Figure 2B). Brain regions downstream of APCx may be able to interpret the total intensity of global APCx output (Figure 5G) as modulated by respiration to make the necessary perceptual decision. However, it is also possible that the downstream decision-making brain regions could be informed largely by those neurons with a particular functional group of neurons – for example, those with strong respiratory modulation (Figure 5H-I, left) but not by those APCx neurons whose responses are independent of the respiratory phase (Figure 5H-I, right). If this were the case, the different groups of neurons identified from the clustering of the phase responses (Figure 5H) will differ in their decoding accuracies in predicting either trial type or behavioral choice. This prediction was confirmed: when task-related variables were predicted based on spikes from neurons belonging to the cluster with maximal modulation by respiratory phase, logistic regression scores were significantly higher than for neurons with lower tuning to the arrival time of the input during the respiratory phase (Supplementary Figure 5A-C).

In summary, the responses of APCx population triggered by odor pulses showed little to no signatures of evidence accumulation. Task-related variables could be represented through small subsets of neurons, with the fidelity of the encoding being affected by the phase-tuning and firing pattern of the individual neurons.

## Discussion

Using a novel behavioral task we have demonstrated that mice can recognize differences in features of fluctuating olfactory stimuli that become apparent only when integrated over several seconds. We also showed that mice valued the odor pulses differentially depending on their timing with respect to the respiratory phase, and that this heterogeneity stems from phase-dependent changes in the amplitude of OSN responses. Modelling of mouse choices revealed that pulse timing relative to breathing phase accounts for some of the errors in choices made by mice. APCx neurons responded stochastically to odor pulses within a trial and their response time courses were heterogeneous. APCx neuron responses were also heterogeneously modulated by the respiratory cycle, suggesting a multiplexed stream of information diverging from highly tuned glomeruli to different neuronal subpopulations in the APCx. While many APCx neurons scaled (up or down) their firing rates with the total odor pulse count, the average firing rates were constant over the duration of the sampling window, rather than ramping up or down over time. This absence of directional changes in firing rates over time with increasing sensory information rules out APCx as a potential olfactory evidence accumulation center. Altogether, our study provides insights into the physiological and behavioral representations of fluctuating stimuli in the context of an olfactory evidence accumulation task.

### The behavioral task

Mice in our task were asked to distinguish and categorize random sequences of brief odor pulses of the same odorant based on whether the integrated signal was below or above a threshold. The signal integrated by mice for decision making could be related to one or a combination of statistical features that characterize odor plumes: intermittency, whiff frequency and concentration profile ^8^. In the current task, the duration of each pulse and the duration of the sampling window were fixed, allowing us to control the values of those statistics by varying the total number of odor pulses delivered. Then, we have shown that when mice were presented with fluctuating olfactory stimuli over several seconds, they could temporally integrate them to make binary behavioral decisions. Our results are not only aligned with recent literature showing that mice are capable of intermittency discrimination ^32^, but also demonstrate that they can successfully distinguish stimuli with minimal variations in intermittency (Supplementary Figure 1C).

The present work also confirms that the temporal integration of olfactory information can occur over time windows covering multiple respiration cycles, in agreement with previous reports using fluctuating stimuli ^28,46^. The random delivery of pulses within the sampling window led to trials with equivalent total pulse counts that differ in their temporal profiles, but more importantly, also allowed trials with pulse counts at both sides of the decision boundary to display similar statistics over short intervals. Good performance in this task requires a decision-making process guided by sensory evidence accumulation, in which mice have to store transient changes in olfactory statistics in a signal that needs to be updated upon encountering newer stimuli. In this context, we observed mice have the flexibility to accurately discriminate stimuli presented in windows ranging from ∼1 to 10 s (Supplementary Figure 1B), similar to what has been reported in rodents for discrimination of visual flashes ^33^ or integration of object quantities in a virtual reality setup ^47^. Whereas sub-second integration times can be detrimental for performance ^48^, there is still an open question about whether evidence accumulation over large timescales is achieved by time-invariant cognitive mechanisms or by flexible adaptation of those mechanisms across different timescales ^49^. For instance, regression analysis on the licking decisions made on 5s trials revealed that mice place different values to olfactory information depending on its timing during the sampling window (Figure 1E). This finding suggests that the storage of early sensory information could be affected by leakage in the integrator, similar to what was previously shown for rats performing multisensory integration for binary choices ^50^.

Sensory evidence accumulation is not only defined by the memory displayed by the accumulator, but also by the level of noise in the accumulating signal. Applying a similar approach to a previous report ^33^, we showed that noise in the estimation of pulses had a constant offset for low pulse counts and then starts to scale with the number of pulses (Figure 2). This observation suggests that the integration of olfactory stimuli follows a subitizing model, in which the noise for the estimation of low quantities is minimal, followed by a monotonically scaling law for higher stimulus counts. However, our noise modelling results challenge subitizing in two main aspects: (i) the estimation of low quantities is not noiseless and (ii) the constant noise extends further than four to five stimuli ^35^. Regarding the first point, noise values were significantly reduced when model parameters were estimated using the perceived odor in a trial -i.e., the odor pulses convolved with the sniff kernel-(Supplementary Figure 2F), suggesting that part of the basal noise derives from perceptual variabilities due to sampling, but the source of the remaining noise offset is still unknown. Regarding the second point, it is important to note that the majority of reports about the limits of subitizing come from human experiments using visual or auditory stimuli, whereas larger limits were found for animal subjects ^34^. Once noise starts to scale with the number of pulses, it is better fitted by a scalar variability model in which the noise scales linearly with the stimulus count. The persistence in the noise scaling after accounting for breathing, suggests that noise scaling is independent of sampling. Scalar variability has been observed in other rodent behaviors involving perceptual evidence accumulation ^33,51^, and it is believed that it can emerge from the neural mechanisms used by the brain to encode quantity estimations ^52^. Although models for counting discrete stimuli seem to be a valid approach to approximate the accumulation of olfactory statistics in our task, the current results and modelling are insufficient to prove that mice are implementing a counting strategy over discretized statistics - such as odor pulse/whiff counts - as opposed to just accumulating temporal variations of a continuous variable -such as odor intermittency or pulse rate. Assays manipulating the duration of the sampling window during catch trials have been used as a strategy to disambiguate the variable that animals integrated ^50^, and they can be incorporated in future experiments.

### Breathing modulation of sensory information

The brief nature of the odor pulses used in our task was essential for testing the effects of breathing in the representation of olfactory stimuli across the ascending olfactory pathway and its later weighing for behavior. Previous reports have shown that MTC responses to steady odor stimuli tile the respiratory phase with a bias towards inhalation ^20,23,53^. Here, we show that OSN responses were larger for pulses arriving during inhalation than for pulses arriving during exhalation, suggesting that OSN activity is contingent on the amount of odor molecules carried by the nasal airflow during breathing, and that breathing-modulated odor responses are already present at the first relay of the ascending olfactory pathway. Our results are related to previous reports on mice discriminating the sub-sniff timing and intensity of brief optogenetic stimulation of OSNs ^24,54^, and confirm that variations in the amplitude of OSN responses elicited by differences in pulse timing within breathing cycle can also be differentially read and integrated by downstream areas for decision-making. MTC responses to odors have been shown to tile the breathing cycle ^20,23,53^, but OSN responses in our experiments have fairly homogeneous breathing phase dependence. This difference suggests that circuits within the OB, likely the abundant and diverse types of inhibitory neurons ^55^, shape the OSN inputs into MTC responses with different phase preferences. Our results also offer a basis for MTCs being able to track changes in the frequency of olfactory stimuli ^22,28,29^. Lastly, there have been descriptions of changes in the amplitude of MTCs responses as a consequence of variations in breathing frequency ^21,26^, which could partly arise from frequency-dependent variation in OSN responses We were unable to test this in our experiments, since mice were familiar with the odorant during the training sessions, there was not much variation in breathing frequency ^56,57^.

The dependence of the amplitude of glomerular responses on the timing of odor pulses relative to breathing can be helpful to understand how olfactory inputs are weighed when animals need to integrate information across multiple breaths. To exclude any effects of sensory adaptation in OSN responses, we focused on changes in GCaMP3 signal for odor pulses that were relatively temporally isolated (time between pulses > 1s). Therefore, how OSNs respond to multiple pulses within a single breath remains to be explored in future work. Experiments in anesthetized, tracheotomized rats have shown that MTC responses to fluctuating odor stimuli scale linearly with stimulus duration, but non-linearly with stimulus concentration ^53^. Moreover, recent work both using calcium imaging and in vivo electrophysiological recordings has suggested that MTC populations across glomeruli seem to be correlated with odor plume temporal dynamics ^30,31^. Recent work tracking OSN responses and behavior to fluctuating odor stimuli have revealed that glomerular responses to different levels of intermittency are heterogeneous, and that heterogeneity is necessary for better predictions of intermittency ^32^. Our results provide functional evidence of the integration of olfactory stimuli over time for odor plume categorization, although complementary experiments are still required to determine which aspect of odor plume dynamics glomeruli track and what the underlying neural mechanisms are.

### Piriform cortex encoding of fluctuating stimuli

We have characterized, for the first time, the response of APCx neurons to highly fluctuating olfactory stimuli. Odors presented passively or within a behavioral task are shown to be represented by not-so-sparse neuronal ensembles in APCx ^58–61^. These studies typically used longer and non-fluctuating odor stimuli and mostly focus on characterizing the size of the ensembles required for reliable odor identity representation. Our results show that when the same odorant is delivered as brief pulses, different subpopulations within the ensemble are recruited each time in a stochastic fashion. In trials in which multiple odor pulses were delivered, timing between the first and second pulse determined the neuronal response to the second pulse, providing clear evidence of sensory adaptation in APCx responses, which could partly account for the variable responses of neurons to each odor pulse. Another source of variability is the differential tuning of APCx neuron response to the timing of odor pulses within the breathing cycle. We found that roughly 50% of the APCx population responded equally to odor pulses falling at any time during the breathing cycle, whereas the remaining 50% may be tuned to different pulse timings over the breathing cycle. It is important to note that neuronal response amplitudes were estimated by the DUNL method by grouping spikes in 50 ms bins, which may influence inferences about the sparseness and phase-tuning of the responses.

What are the signals that shape the phase-tuned odor responses in APCx? One possibility is that a subpopulation of APCx neurons may inherit the tuning of their MTC inputs. Recurrent inhibitory signals in the APCx appear to be essential in controlling the timing and gain of APCx population responses to odors ^59^, and may contribute to phase tuning observed in our experiments. Moreover, a recent report has shown that, even in the absence of odors, inhalation can trigger bidirectional changes in the firing rates of APCx neurons, driven mostly by feedforward bulbar input, but with a small top-down component ^27^. Future experiments can test whether the functional differences we observe are correlated with molecular genetic identity and the connectivity of APCx neurons.

Do olfactory inputs trigger similar response profiles across the APCx neuronal population? The neuron-specific response kernels revealed that the APCx population is also heterogeneous in the time course of their responses to olfactory inputs. It is possible that different kernels may correspond to different cell-types and their connections within the APCx. APCx harbors a molecularly and anatomically diverse population of excitatory cells ^41,62^, and also two subpopulations of inhibitory cells which have been shown to engage in local feed-forward and feed-back inhibitory circuits ^59^. In fact, Bolding and Franks (2018) described how those inhibitory circuits were essential for silencing APCx population responses after a peak firing during the first sniff. Interestingly, we did not find a correspondence between the clusters defined by the kernels and those defined by the phase-tuning, suggesting that APCx neurons with a specific phase-tuning can diverge in terms of their response profile and vice-versa. However, we take the kernel-based clusters as a global snapshot of the functional diversity within the APCx, and their correspondence with neuronal clusters defined by gene-expression or projection patterns will need to be clarified with future experiments.

### Sensory vs. decision information in piriform cortex

We found that task-related variables could be decoded from a small number of APCx neurons (∼20), if the entire sampling window is considered. This might not be surprising if we take into account that subpopulations of ∼100 randomly sampled APCx could accurately classify different odor identities ^58,60,63^. In those previous reports as well as in the current work, we found that further increasing the number of neurons did not improve classification accuracy, supporting the idea of high redundancy in the encoding of olfactory inputs at the APCx. Higher classification accuracies were found for trial identity compared to choice, providing the first indication that APCx population firing might be reflecting the time course of sensory inputs, rather than any sort of neural correlate of evidence accumulation. This idea gets additional support after finding the classification accuracy for APCx neuronal firing does not significantly increase over time for independent time intervals spanning the sampling window. Moreover, previous studies in other sensory modalities have shown that despite reliably responding to sensory inputs, and the impairments in performance after their inactivation, early sensory cortices may not be responsible for sensory evidence accumulation ^64–67^. The fact that the amount of sensory input is unbalanced between the two trial types poses an additional caveat of the interpretation of these results. In the future, perturbation experiments, as well as behavioral assays in which the integrated olfactory signal could be balanced between the two trial types in the current task - for instance, computing the ratio of pulses between two different odorants - will provide further clarification about the role of APCx in the encoding of olfactory statistics. Different clusters of APCx neurons achieved different decoding accuracies, but whether the most accurate neurons have specific projection patterns to non-olfactory areas that might be involved in decision making remains to be seen.

Our study leaves open the question of where in the mouse brain olfactory statistics are accumulated for decision-making during the task. APCx sends both extensive projections to other olfactory areas, and also direct projections to the orbitofrontal cortex, medial prefrontal cortex, and medial dorsal thalamus ^68^. Extensive work in mice assessing accumulation of visual or auditory evidence have identified areas in the parietal and prefrontal cortex responsible for the graded accumulation of sensory inputs and the consequent categorization of the accumulated value into a behavioral decision ^43,66,67,69^. It might be likely, then, that olfactory evidence could be also accumulated in those areas. Additionally, recent rodent work has shown the involvement of the anterior dorsal striatum in auditory evidence accumulation ^70^, a relevant finding if we consider that olfactory inputs can reach the ventral striatum through the olfactory tubercle ^71–73^, an area associated with the encoding of odor valence, but with direct connections from OB and APCx. If the temporal integration or updating of olfactory stimuli plays a role in naturalistic behaviors such as trail following, which may also involve multimodal integration, it is highly likely that similar brain hubs for sensory evidence accumulation are involved for the different sensory modalities.

## Methods

### Experimental animals

All experimental animals used for behavior and electrophysiology were C57Bl6/J mice of either sex acquired from Jackson Laboratories, aged two to four months at the start of experiments. Following the implantation of a tetrode drive and/or head plate, all mice were housed individually. Behavior and physiology experiments took place over the course of one to two months. After the completion of all experiments, mice were euthanized, and post-mortem histology was performed to confirm the location of electrophysiological recordings. Adult heterozygous OMP-GCaMP3 ^74^ from a breeding stock maintained within Harvard University’s Biological Research Infrastructure were used for the olfactory bulb imaging. All mice used in this study were housed in an inverted 12 h light cycle and fed ad libitum. Animals were housed at 22 ± 1 °C at 30–70% humidity. All the experiments were performed following the guidelines set by the National Institutes of Health and approved by the Institutional Animal Care and Use Committee at Harvard University.

### Behavioral apparatus

A custom-made apparatus was built inside a Faraday cage to allow the execution of behavioral experiments simultaneously with electrophysiology and imaging. Animals were head-fixed over a styrofoam ball which allowed them to freely walk during behavioral sessions. A pair of lick ports were appropriately positioned in front of the animal through a 3D printed platform coupled to a stepping motor controlled from a TinyG v8 microcontroller (Synthetos). Water delivery was executed from an Arduino-Mega (Arduino), controlling two 3-way solenoid valves (LHDA12333115H, Lee Company). Licking detection was achieved through a capacitive circuit.

Odor pulse delivery was accomplished using a custom olfactometer equipped with fast proportional solenoid valves (EV-P-05-09-A0; Clippard) controlled from a Teensy 3.2 (PJRC). The fast kinetics of the solenoid valves used allowed us to switch between constant clean air delivery and 50 ms activations of another solenoid valve connected to a vial containing 5% (v/v) ethyl valerate (Sigma-Aldrich), keeping a constant airflow of 1L/min and without introducing pressure changes that could potentially be sensed by the animal. Then, the resulting odor stream was sent directly to a 3D printed mask fitted to the snout of the animal. For behavior and electrophysiology experiments, ethyl valerate dilutions were done in mineral oil (Sigma-Aldrich), whereas for imaging experiments they were done using di-ethyl-phthalate as solvent (Sigma-Aldrich). The mask was also connected to an airflow sensor (AWM3100V; Honeywell) to monitor animal breathing during the task, as well as to a vacuum line to ensure quick removal of the odorized air from the snout of the mouse. Vacuum flow from the mask, as well as the airflow input to the olfactometer, were regulated using two different mass-flow controllers (QPV1, Proportion Air). Photoionization detector (PID; Aurora Scientific) measurements were performed in absence of any animals but in the same conditions of an experimental session, with the probe of the PID placed at the approximate position of the nostrils of the mouse.

Global task structure and communication with the different microcontrollers and cameras was achieved using custom-written software in Python. Behavioral data (licking and breathing) was acquired using a PCIe-6351 card (National Instruments).

### Behavioral task

Task structure was controlled by custom-written software in Python. Each trial in the Poisson-distributed Odor Pulses Task comprised three different periods (Figure 1). The first one – “Waiting Period” – had a 5s duration and its ending was marked by a 10 kHz sound cue. The second period - “Sampling Period”- could have different durations across different training and testing sessions (see Behavioral Training below) and was the time window during which odor pulses were delivered and lick ports were moved away from the animal by a motor. A 6 kHz sound cue indicated the end of the “Sampling Period” and the displacement of the lick ports closer to the animal, leading to a 1.5 s “Response Period” in which the animal had to report its decision by licking one of the lick ports according to the total pulse count. If the animal made the correct licking choice, a 7 µL water drop was delivered, whereas if the lick was in the wrong lick port no water was delivered, a 3.8 kHz buzz tone was presented, and a 10 s timeout was given. A regular behavioral session consisted of 100-200 trials.

### Behavioral training

One week after surgery, mice were habituated to be handled and head-fixed by experimenters. Next, animals were put under water restriction in compliance with approved protocols. On the first day of water restriction, mice were given 20 minutes of free exploration of the behavioral apparatus with water available from lick ports. In the next session, animals were head-fixed in the dark, and water drops were delivered frequently to encourage them to find the lick ports. Then, animals were moved to the ‘Lick Training’ phase, in which they were presented for the first time with the trial structure, but without delivering any odor pulses. During this phase, water was automatically delivered to either of the two lick ports after the second sound cue to encourage animals to lick from the lick ports to get the reward. Lick port location adjustments and manual delivery of water to neglected lick ports were done -if needed- to avoid the development of biases towards a specific side. Once animals showed commitment to lick until they were satiated, they were moved to the ‘Block training’ phase in which the actual task structure was introduced, with odor pulses being delivered during a 1 s sampling window. At this point, the total number of pulses in a trial could be drawn from two different Poisson distributions with different mean values: *λ*_1_=16 for ‘High Trials’ and *λ*_2_=1 for ‘Low Trials’, with both types of trials presented in blocks of 17 trials. Also, for the first 1-2 sessions of ‘Block Training’, water was delivered automatically to the rewarded side to help the animal to learn the association between the number of pulses and the side of the reward. After the second session in this phase, water was only delivered upon the animal licking in the side associated with reward for that type of trial. Block size was progressively reduced after animals reached more than 80% success in a session until the two types of trials were delivered randomly. Then, keeping the random trial structure, the difference between *λ*_1_and *λ*_2_ was progressively reduced as well as the sampling window duration was increased. The whole training process took ∼2.5 weeks.

### Olfactory Bulb Craniotomy

A craniotomy was performed to provide optical access to both olfactory bulbs. Mice were first anesthetized with an intraperitoneal injection of ketamine and xylazine (160 and 16 mg/kg, respectively) and the eyes were covered with petroleum jelly to keep them lubricated. Before starting the surgery, carprofen (7.5 mg/kg) and dexamethasone (10 mg/kg) were injected subcutaneously, followed by an intramuscular injection of cefazolin (500 mg/kg). Body temperature was maintained at 37 °C by a heating pad. The scalp was shaved and then opened with a scalpel blade. After thorough cleaning and drying, the exposed skull was gently scratched with a blade, and a titanium custom-made headplate was glued on the scratches with Loctite 404 Quick Set Adhesive. The cranial bones over the OBs were then removed using a 3 mm diameter biopsy punch (Integra Miltex) and then the brain surface was cleared of debris. The exposed brain area was kept moist with artificial cerebrospinal fluid containing (in mM) 125 NaCl, 5 KCl, 10 Glucose, 10 HEPES, 2 CaCl2 and 2 MgSO4 [pH 7.4], as well as Gelfoam soaked in the same solution (Patterson Veterinary). Two 3 mm No. 0 glass coverslips (Warner) were glued together with optical adhesive (Norland Optical Adhesive 61) and adhered to the edges of the vacated cavity in the skull with Vetbond (3M). C&B-Metabond dental cement (Parkell, Inc.) was used to cover the headplate and form a well around the cranial window. After surgery, mice were injected with slow-release buprenorphine (3.25 mg/kg). Animals were allowed to recover for at least 7 days.

### Wide-Field Calcium Imaging

Wide-field imaging was performed on adult OMP-GCaMP3 mice as described previously ^75^ using a 4X objective (Olympus, NA 0.1). Blue light from an LED (M470L3, Thorlabs) was used for excitation, and the emitted light was filtered (MF525-39, Thorlabs) and collected with a CMOS camera (BFLY-U3-23S6M-C, FLIR). Videos were acquired using a 10 Hz frame rate and a 2×2 binning of the image.

### Chronic Tetrode Implantation

Mice were anesthetized with an intraperitoneal injection of ketamine and xylazine (100 mg/kg and 10 mg/kg, respectively), then placed in a stereotaxic apparatus. After the skull was cleaned and gently scratched, a custom-made titanium head bar was glued to it. A small craniotomy was performed above the implantation site, before 8 custom-built tetrodes (Chang et al., 2013) were lowered together into the brain (coordinates for APC: antero-posterior 1.6mm, medio-lateral -2.8mm, dorso-ventral 3.4mm; all antero-posterior and medio-lateral coordinates are given relative to Bregma; all dorso-ventral coordinates are given relative to the brain surface). A reference electrode was implanted on the occipital crest. All the mice were implanted in the right hemisphere. The whole system was stabilized with dental cement. Mice were given a week of recovery before any new manipulation.

### Neuronal Recordings

A week after chronic tetrode implantation, mice were habituated and trained in the task as described above. Brain activity was recorded once a day, always at the same period of the day and only once mice reached proficiency in the task. Recordings were conducted while animals were awake and head-restrained, and data was only collected during the 5 seconds ‘Sampling Window’ period of each trial. However, the first 500 ms of the recordings were discarded due to artifacts. Since the headstage allowed tetrodes to be finely adjusted up or down, tetrodes were slightly lowered in the brain after each recording session (around 40µm deep), ensuring that different neurons were recorded each day. Electrical activity was amplified, filtered (0.3-6kHz), and digitized at 30kHz (Intan Technologies, RHD2132, connected to an Open Ephys board). Single units were sorted offline manually using Kilosort v1 ^76^. Units with more than 1% of their inter-spike intervals below 2ms refractory period were discarded. Units displaying large changes of amplitude or waveform during the recording were also discarded ^77^. The position of the tetrodes in the brain was confirmed post-mortem through electrolesion (200μA for 4s per channel).

### Noise Modelling

The modeling of how the total number of pulses delivered, *N_p_*, contributes to the decision noise was inspired by a previous report ^33^. Our model assumed that the decision-making process of a mouse in each trial starts with the estimation of the total number of pulses delivered (*N*_est_), followed by the comparison of this value to the estimation of the threshold for trial categorization (*N_b_*).

As is shown in Eq. 1, we assumed *N_est_* followed a normal distribution centered around *N_p_*, with a variance *σ^2^* , which constitutes the first source of decision noise and reflects the noise in the estimation of the total number of pulses.

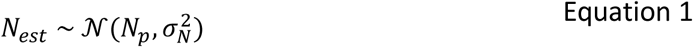

The second source of noise arises from the internal estimation of the *N_b_*, a parameter that we also assumed follows a normal distribution with constant variance *σ^2^* for all *N* . T Equation 2 shown in Equation 2, we propose that licking decision on a trial is based on ΔN, i.e. the internal representation that the animal has of the difference between sensed pulses (*N_est_*) and the decision boundary (*N_b_*).

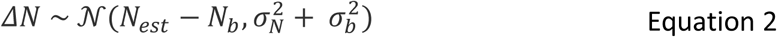

Therefore, the probability of mouse choosing the ‘High’ side is equal to the probability of ΔN being greater than 0, which corresponds to the area under the Gaussian curve defined by Equation 2:

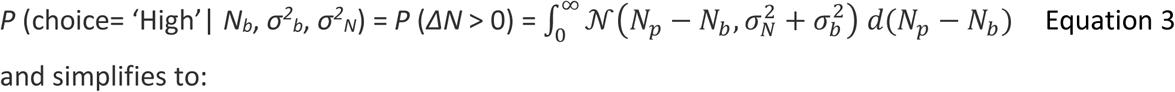

and simplifies to:

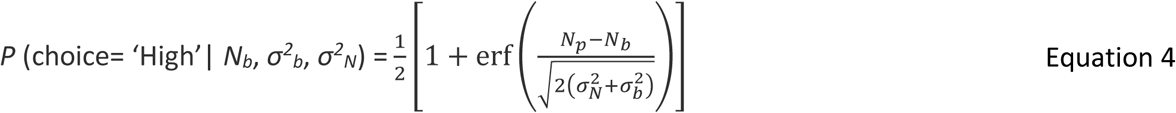

Fitting the model with the values of Np and the mouse choice in each trial, we can obtain the maximum likelihood estimation of the parameters. Confidence intervals of parameters were estimated through bootstrapping 20000 trials with replacement over 1000 iterations.

The likelihood function can be described with a total of 22 parameters (*N_b_*, *σ_b_* and 20 *σ_N_*‘s corresponding to distinct numbers of pulses). Note however, that since the variances appear as 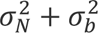 we will define 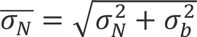 and find these values instead. Further, note that *N_b_* should correspond to the point where the psychometric probability crosses 0.5. However, because of the discrete nature of *N_p_*’s, *N_b_* could be determined up to an interval between two integers. We choose *N_b_* to be the closest integer to the point of 0.5-crossing, which in our data leads to *N_b_* = 8. This leaves us with 20 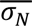 parameters to be calculated from MLE.

For testing the different models of noise scaling to the behavioral data, maximum likelihood estimation of the specific parameters of each of the models -and their correspondent confidence intervals-was performed bootstrapping 15000 trials with replacement over 1000 iterations.

### Mixed-models Logistic regression

Odor pulses in every trial were convolved with a kernel derived from PID recordings, and odor signal was binned and averaged for every bin. Then, the contribution of odor information to mouse’s licking decision was assessed by fitting two different mixed effects logistic regression models using Pymer4 package ^78^. The first model was fitted specifying a single, shared coefficient for all bins of odor information, whereas the second model was fitted allowing each bin of odor information to have different coefficients. Therefore, the models represent scenarios in which odor information is weighed equally or differentially across the whole ‘Sampling Period’. Models were compared using a log-likelihood ratio test.

### Analysis of APCx spiking data

For each behavioral session, spikes from the entire APCx neural population were used to fit a linear regression model aimed at predicting the total number of odor pulses delivered on a trial. Then, only those neurons whose spikes led to a reduction in the Akaike Information Criteria (AIC) of the regression model fitted for every session were considered for all subsequent analysis (except for the dependence on the decoding with the size of population in Figure 4 F-H). For each session, a linear regression model was fitted to predict total odor pulse counts based on the spiking of APCx population, whereas logistic regression models were used to predict variables with binary outcomes, such as trial identity (‘High’ or ‘Low’) or mouse choice (‘High side’ or ‘Low side’). To test whether APCx neurons were accumulating evidence for decision-making, APCx firing during odor pulse presentation was divided into different terminals, and regression models were ran using the spiking in cumulative or independent intervals.

#### Analysis of APCx neural responses using deconvolutional unrolled neural learning (DUNL)

Single-trial neural activity was decomposed into impulse-like responses to odor pulse presentation as it has been described in a previous report ^45^. Briefly, the mean activity of a given neuron in a trial was expressed as the convolution of a kernel -characterizing the response of the neuron to an odor pulse-with sparse codes - a vector representing the timing of events, together with the strength of the neural responses associated with those events. Kernels were learned fully from data, and in this dataset a single kernel was learnt for each APCx neuron. Non-zero elements of the sparse code were aligned to the timing of the odor pulses. Stochasticity in the estimated activity was added by passing the convolved signal through a generative model using a Poisson process. For convenience, we are renaming the code amplitudes as ‘response amplitudes’ in Figure 4. Analysis on response amplitudes and kernels from APCx neurons was only performed in those neurons that were considered for the analysis of the spiking data. ‘Phase clusters’ were obtained by performing K-means clustering on the response amplitudes elicited by pulses at different times during respiratory phase in each neuron. ‘Kernel clusters’ were obtained after performing K-means clustering on the 1s kernels learned for each neuron.

#### Analysis of imaging data

Images were processed using both custom and available MATLAB (MathWorks) scripts. Motion artifact compensation and denoising were done using NoRMCorre^79^. The CaIMaN CNMF pipeline ^80^ was used to select and demix ROIs. dF/F values were calculated using the mean of a baseline period of at least 20 frames. To identify glomeruli that were significantly odorant-modulated, three points centered on the peak dF/F signal after odorant delivery were averaged. Glomeruli were classified as significantly odorant-modulated if their peak averaged response exceeded 2.5 standard deviations of the baseline noise.

## Supplementary Figures

**Supplementary Figure 1.**
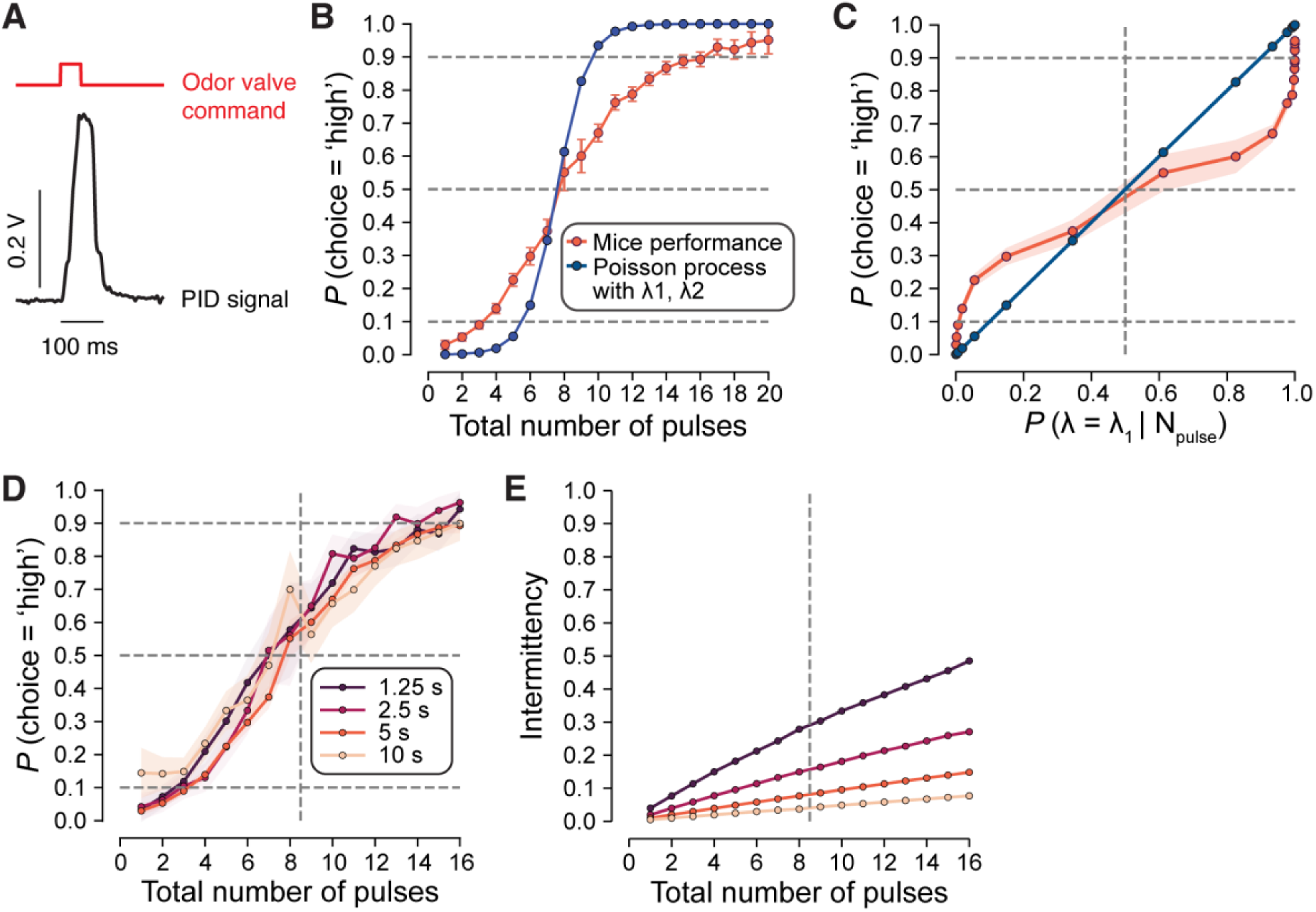
**(A)** PID recordings (black trace) after the delivery of a single 50 ms pulse of ethyl valerate 5% (red trace, voltage command to the solenoid). Temporal scale is the same for both traces. An estimation of the odor signal in each trial could be now generated by convolving the temporal course of voltage command to odor valves with this PID profile obtained from a single pulse. **(B)** Comparison of the psychometric curve obtained from mice choices (orange trace, same as the one in the bottom panel of Figure 1D) and the psychometric resulting from calculating the probability of binary choice assuming the animal is making the choice based on the ratio of conditional probabilities based on the known values of λ_1_ and λ_2_ for the two Poisson distributions generating the total pulse count on each trial (blue trace). The λ_1_ and λ_2_ values used were the same as those used for generating total pulse counts in a trial for the behavioral data. **(C)** Similar to B, the probability of choosing the ‘High’ side was computed as a function of the probabilities calculated based on the ratio of probabilities between the two Poisson distributions used for generating the total pulse counts on the trial. Mice choices does not follow the choice predictions based on the ratio of probabilities arising from the two Poisson distributions. **(D)** Psychometric curves obtained with a λ_1_/λ_2_ = 3 but using different sampling window durations. The curve corresponding to the 5 s sampling window is the same as shown in Figure 1D. Sample sizes: 1.25 s (4 animals, 2760 trials), 2.5 s (4 animals, 2533 trials), 5 s (7 animals, 20042 trials), 10 s (4 animals, 2715 trials). **(E)** Intermittency of the odor signal for all total pulse counts in the conditions described in D. For B-E, error bars represent 95% confidence interval.

**Supplementary Figure 2.**
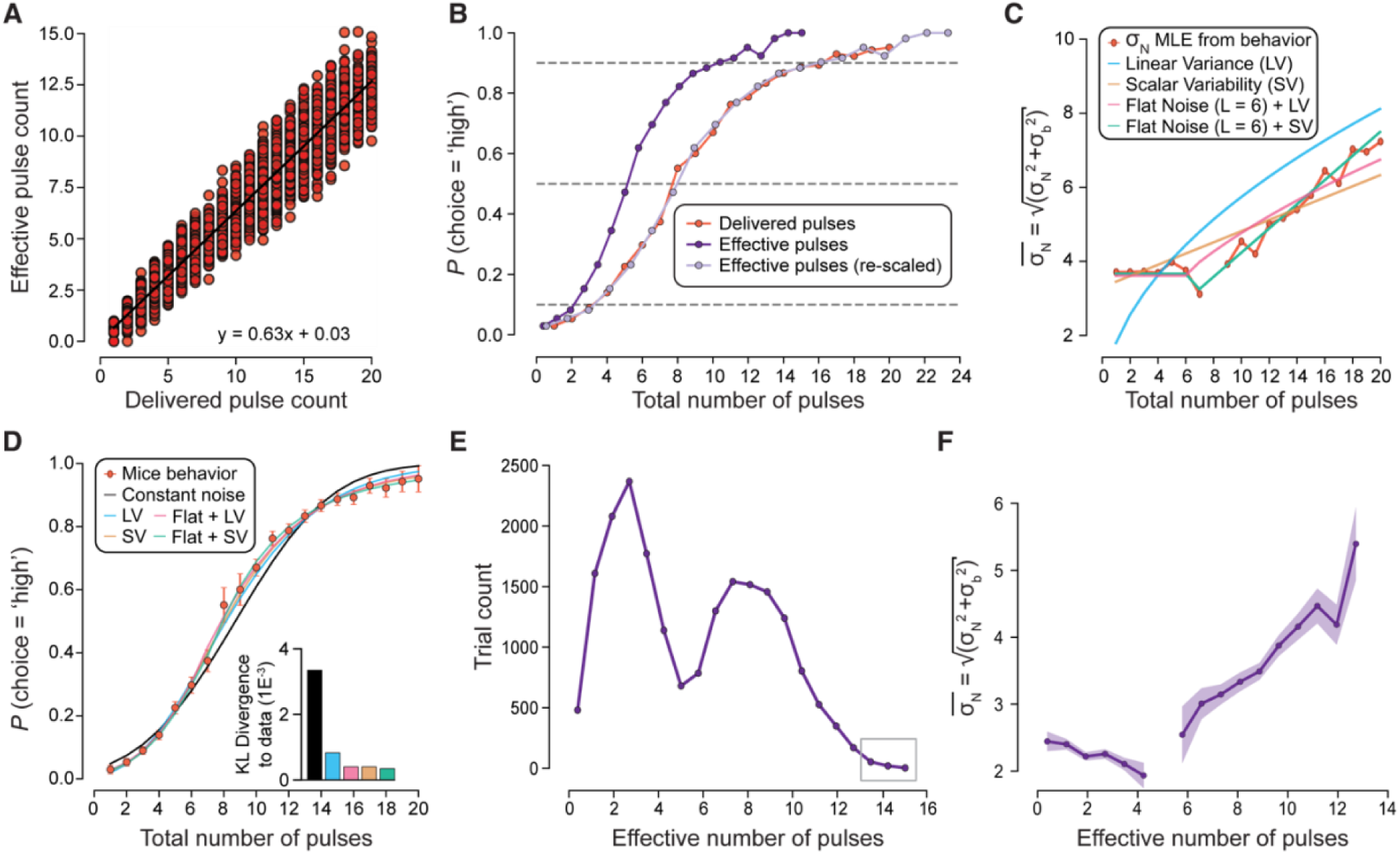
(**A**) Regression between the delivered pulse counts in each trial and their corresponding effective pulse counts calculated by the linear combination of the phase histograms and the coefficients from B. **(B)** Comparison of the psychometric curve based in the total delivered pulse count (orange trace, same as in Figure 1D), the psychometric based in the effective pulse count -i.e. after convolving the phase histograms of each trial with the logistic regression coefficients from Figure 2B-(dark purple trace), and the psychometric curve based on effective pulse counts after re-scaling based on the linear regression shown in Figure 2C (light purple trace). **(C)** MLE estimates of 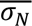 (same as in Figure 2E) and the predicted 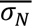 values according to the different models of noise scaling. Scalar variability (SV) models assume a linear scaling of noise with stimuli count, whereas linear variance (LV) models assume a square root scaling of noise with stimulus count. Flat noise models assume constant noise values for low pulse quantities until a limit, followed by some form of noise scaling -such as LV or SV. **(D)** Fits of different models of noise scaling with stimulus number to the observed psychometric curve (orange trace, same as in Figure 1D). For comparison, we also show fits of a model assuming a constant value of 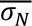 irrespective of the total number of stimuli presented (‘Constant Noise’). **Inset**: Kullback-Leibler (KL) divergence to data was computed for all models. The model with the best fit to data was the one assuming a flat noise followed by a linear scaling of noise with the number of stimuli (Flat Noise (L=6) + SV). **(E)** Histogram of number of trials across the whole range of effective pulse quantities. Since bins within the gray box had very low occurrence, they were pooled together with the closest bin (bin center = 12.72) for MLE calculations. **(F)** MLE estimation of 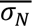 across effective pulse count values without rescaling.

**Supplementary Figure 3.**
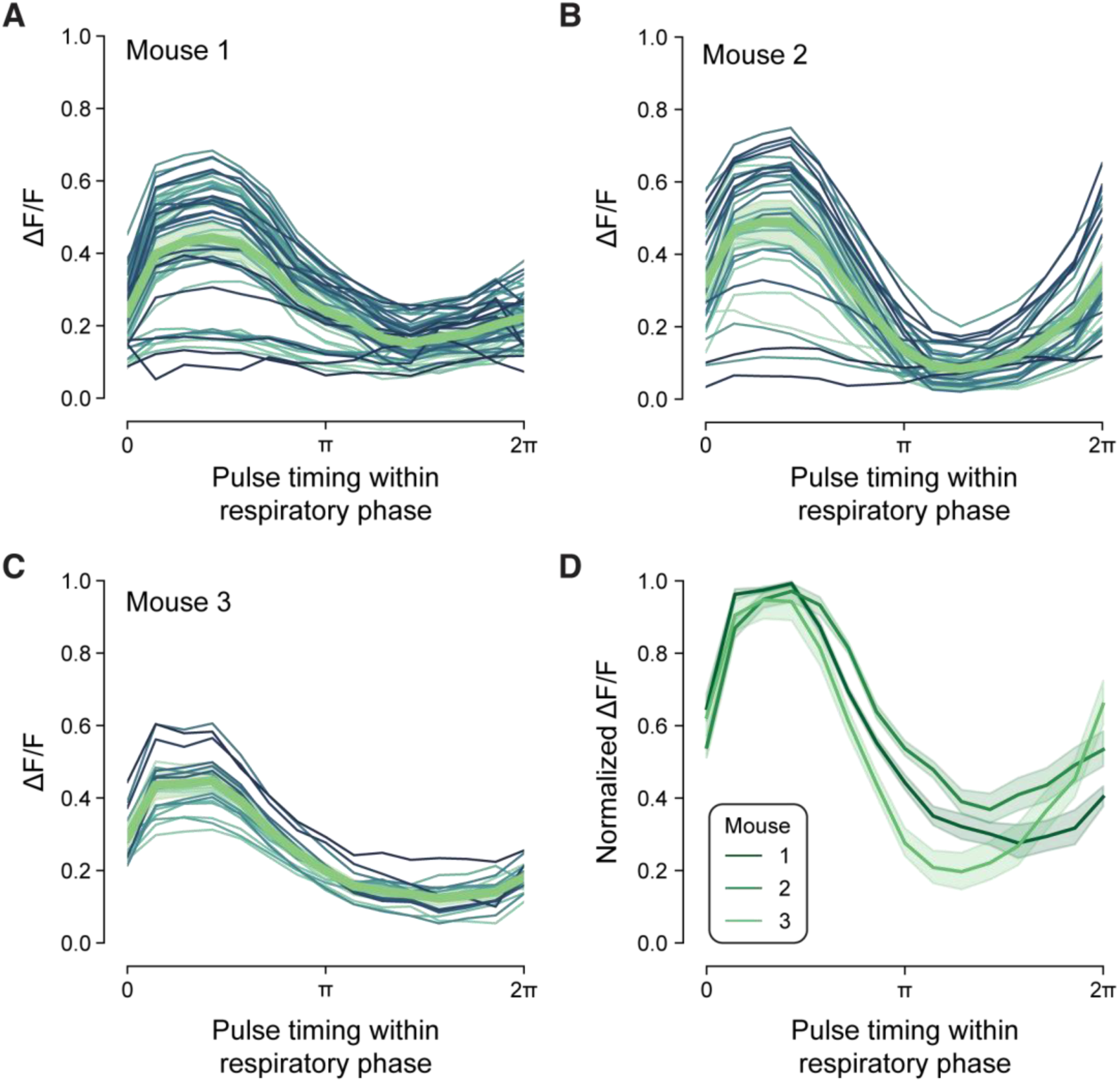
(**A-C**) GCaMP3 fluorescence (ΔF/F) as a function of odor pulse timing relative to the respiratory phase for the three different animals tested. Each thin trace corresponds to a single glomerular ROI (all ROIs that showed significant GCaMP3 response), whereas the thick green line is the average across ROIs. The dependence on pulse timing for response amplitude can be verified even at the level of single glomerulus. (**D**) Normalized GCaMP3 ΔF/F as a function of odor pulse timing relative to the respiratory phase averaged across all the ROIs in each animal. Data in this panel is expressed as mean ± CI 95% (n = 3 animals).

**Supplementary Figure 4.**
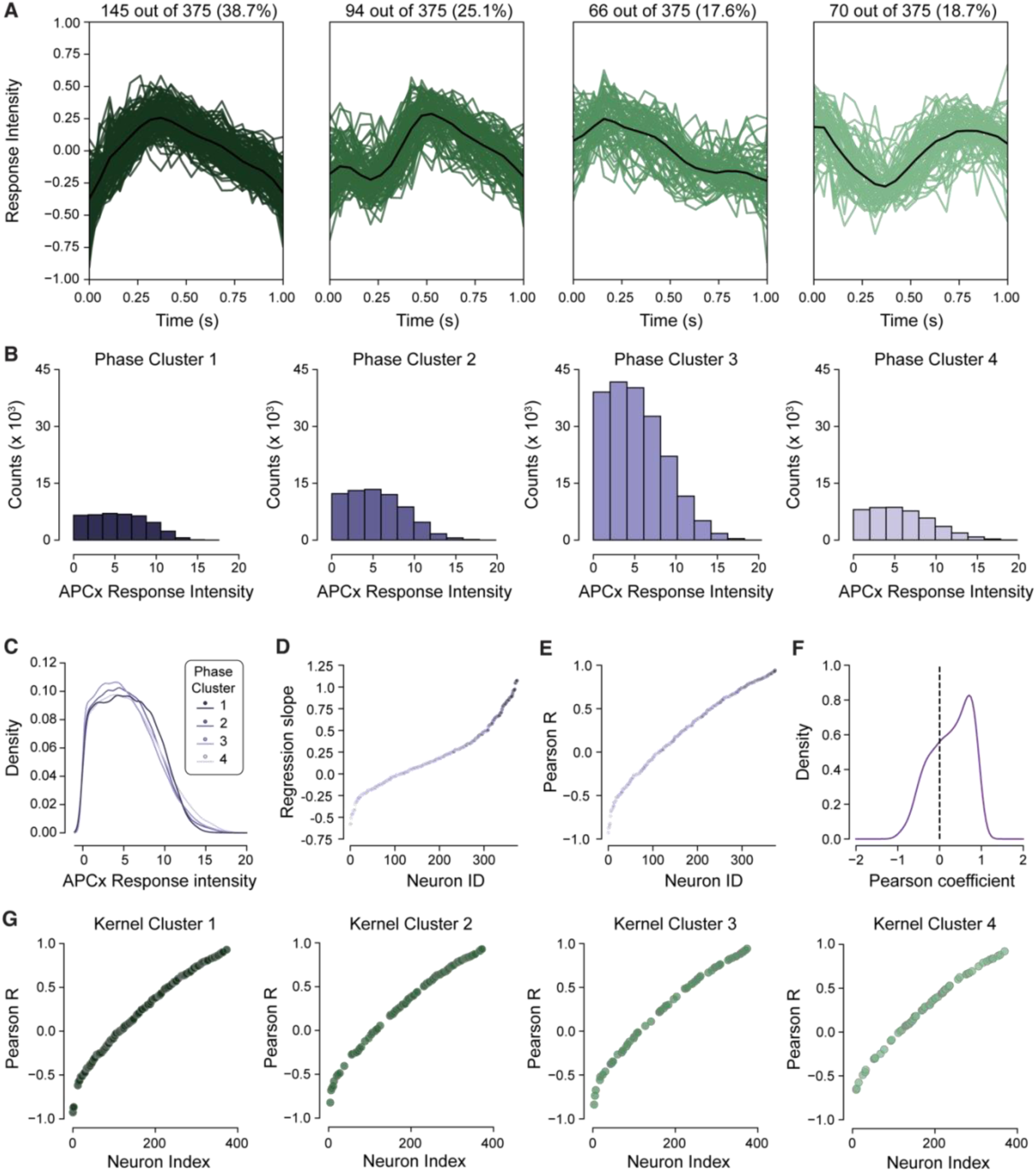
**(A)** Four clusters identified by K-means clustering of the zero-mean kernels obtained for every neuron from DUNL. Colored lines represent individual neurons, whereas the black line indicates the average for the cluster. **(B)** Histogram of APCx response intensities for each of the phase clusters described in Figure 5H-I. **(C)** Kernel Density Estimations of the APCx responses for each cluster. Jensen-Shannon divergences computed across pairs of phase cluster densities were all below 0.0072, suggesting that distributions were all nearly identical. **(D)** Sorted slopes of the linear regression between the response of every APCx neuron to pulses arriving at different times during the respiratory phase and the mean normalized OSN phase responses obtained from calcium imaging. APCx neurons were sorted based on the slope of the linear regression and colored based on the cluster that were assigned from K-means clustering of the respiratory phase modulation of pulse-triggered neuronal responses. Responses from neurons that clustered together tend to share similar levels of correlation with the OSN responses **(E)** Same as B, but for the Pearson coefficients of the correlation between the response of every APCx neuron and the mean OSN phase response from Figure 3C. Similar to what we observed for the slopes, the distribution of Pearson coefficient values suggested a gradient in tuning across the recorded APCx population **(F)** Kernel Density Estimation of the distribution of correlation coefficients in C. The distribution did not follow a normal distribution (Kolmogorov-Smirnov test, p-value = 1.21 E-23) and was skewed-left, supporting the existence of a significant group of neurons with high respiratory-tuning in their sensory-triggered responses **(G)** Neurons sorted using a similar approach as in C, but separating and coloring the neurons based on the clusters obtained from K-means clustering of the neuronal kernels. In this case, neurons that clustered together based on their kernels did not necessarily match in terms of the correlation between their phase responses and OSN activity.

**Supplementary Figure 5.**
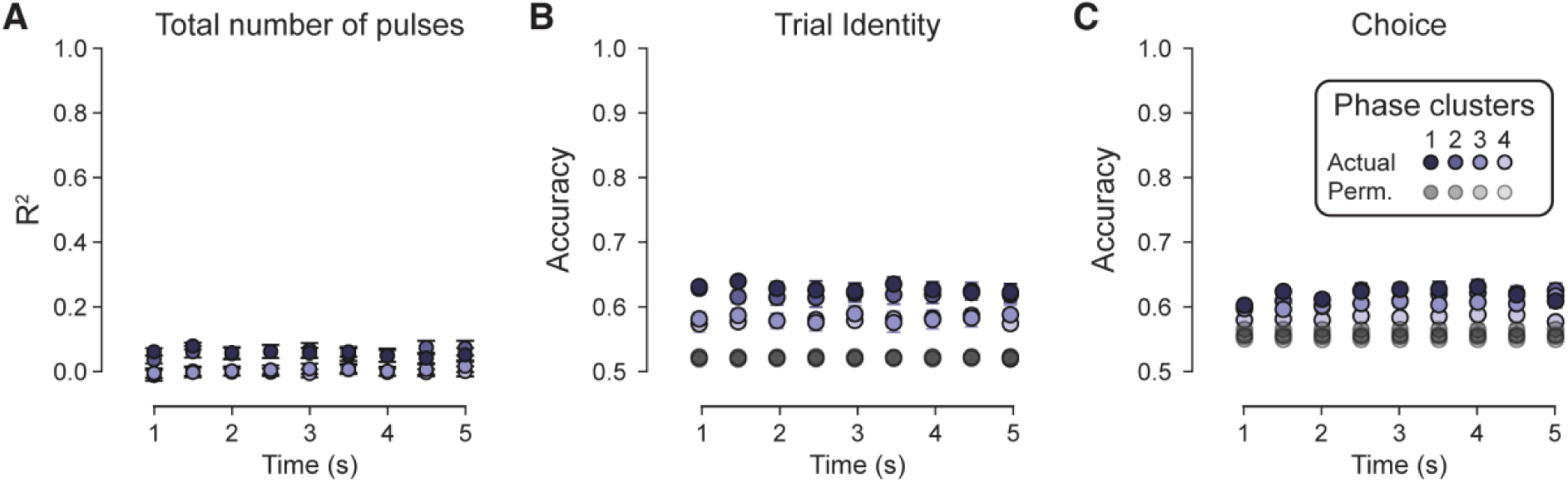
**(A-C)** Same analysis as in **Figure 4 C-E** but running the regressions separately for the spikes over 500 ms independent time intervals of neurons in each of the 4 clusters obtained from K-means clustering of the APCx phase responses. Data is shown as mean ± CI 95%, n = 3 animals over multiple recordings sessions.

## Author contributions

V.N.M., L.E.B. and H.W. designed the project and V.N.M. supervised it. L.E.B. and H.W collected and analyzed the data. H.W., P.M., F.P., adapted and refined the signal-detection theory model for the analysis of decision noise. J.Z. analyzed the calcium imaging data. B.T. and D.B. developed and applied DUNL algorithm to the data. S.J. assisted with data collection. L.E.B and V.N.M. wrote the manuscript, with edits from all other authors.

## Acknowledgments

We thank Julien Grimaud for assistance with the surgeries for *in vivo* recordings. We also thank Jacob Zavatone-Veth for suggestions and comments. We also thank Ed Soucy and Yuwei Li from Harvard FAS Neuroengineering Core for technical assistance. This work was partially supported by startup funds from Harvard University and a grant from NTT Research to V.N.M. L.E.B was also supported by Pew Latin American Fellows Program. F.P was supported by the CBS-NTT Research Physics of Intelligence program.

## Notes

### Competing Interest Statement

The authors have declared no competing interest.

